# The hidden effects of agrochemicals on plant metabolism and root-associated microorganisms

**DOI:** 10.1101/2021.03.14.435313

**Authors:** S. Cesco, L. Lucini, B. Miras-Moreno, L. Borruso, T. Mimmo, Y. Pii, E. Puglisi, G. Spini, E. Taskin, R. Tiziani, M. S. Zangrillo, M. Trevisan

## Abstract

Agrochemicals are commonly used in agriculture to protect crops and ensure yields. Several of them are mobile within the plant and, being perceived as xenobiotics regardless of their protective/curative roles, they induce a reprogramming of secondary metabolism linked to the detoxification processes even in the absence of phenotype symptoms. Moreover, it is well documented that plants, thanks to the root exudation of different metabolites, are able to shape the microbial population at the rhizosphere and to significantly affect the processes occurring therein. Here we show that plant metabolic response to foliarly-applied pesticides is much broader than what previously thought and includes diverse and compound-specific hidden processes. Among others, stress-related metabolism and phytohormones profile underwent a considerable reorganization. Moreover, a distinctive microbial rearrangement of the rhizosphere was recorded following foliar application of pesticides. Such effects have unavoidably energetic and metabolic costs for the plant paving the way to both positive and negative aspects. The understanding of these effects is crucial for an increasingly sustainable use of pesticides in agriculture.

**Highlight:** The foliar application of pesticides induces a broad metabolic reprogramming in plant and shapes the microbial population of the rhizosphere.

## Introduction

It is generally recognized that agronomic practices must include precise control of pests (fungi and insects) infection and weeds invasion, to ensure an economically sustainable harvest. These agrochemicals are perceived as xenobiotics by the plant. To date, a large amount of research has been produced regarding pesticide residues and fate into the environment. Literature indicates that crops are known to be able to take up, translocate, and metabolize xenobiotics in an attempt to detoxify them. Nonetheless, increasing evidence is suggesting that xenobiotics may disrupt and alter plant functions and processes and cause oxidative stress even in the absence of stress symptoms (Parween *et al*., 2016; Shahzad *et al*., 2018). In turn, considering that plants can release up to 25% of the carbon assimilated by photosynthesis in the rhizosphere (*i*.*e*., the soil-microorganism-root interface), such metabolic perturbations likely affect quali-quantitatively the pool of metabolites exudated by roots. As a consequence, the microbial community colonizing the rhizosphere is likely reshaped, with implications on plant growth and soil fertility (Mimmo *et al*., 2014). The comprehension of such hidden perturbations can be of crucial role in the framework of a holistic environmental perspective, in order to reconsider this agronomic practice in the context of increasingly sustainable agriculture (Quintero-Angel and González-Acevedo, 2018). To this aim, the present work was focused on elucidating the biochemical perturbations underlying the use of pesticides, together with the resulting changes at the root microbiota level. To this aim, tomato (*Lycopersicon esculentum* Mill.) has been used as a model crop because of its broad and diverse secondary metabolism. The fungicide tebuconazole, the insecticide imidacloprid, and the fertilizer urea were selected as representative agrochemicals and applied as recommended in the labels by the manufacturers. These agrochemicals, all authorized for use on tomato, were foliar applied in agreement to common agricultural practice. An additional treatment, using the non-selective herbicide glyphosate, was intentionally included as a reference where metabolic disruption was beyond doubts (positive control). Plant and rhizosphere samples were collected after 2 and 4 days from each treatment. High-resolution mass spectrometric metabolomics was applied to disentangle plant biochemical perturbations, and high throughput sequencing of phylogenetic markers for both bacteria (16S amplicons) and fungi (ITS) applied to profile the microbial communities in the rhizosphere.

## Materials and methods

### Reagents

Tomato (*Solanum lycopersicum* L.) cv. Marmande seeds were purchased from SeedsSelect (Fuscello Agostino Sementi srl., Andria BT, Puglia, IT). The fungicide Folicur (tebuconazole 300 g L^-1^ SC), the insecticide Confidor (imidacloprod 300 g L^-1^ EC), and the herbicide Roundup (glyphosate 300 g L^-1^ SC) were commercial formulations bought from Bayer Cropscience, Milan Italy. The fertilizer urea was purchased from Biuron (Borealis Agrolinz Melamine GmbH, Linz, AT). Water for the control treatment consisted of grade II ultrapure deionized water (HOH Water Technology, Palatine, USA). Formic acid, methanol and water used in the metabolomics experiments were LCMS grade from Merck.Metagenomic experiments used the DNeasy PowerSoil Kit (QIAGEN GmbH, Hilden, Germany), Phusion Flash High-Fidelity Master Mix (Thermo Scientific Inc, Waltham, MA, U.S.A.), PCR ultrapure nuclease free water (Thermo Scientific Inc, Waltham, MA, U.S.A.), and Agencourt AMPure XP (Beckman Coulter, Milan, Italy).

### Plants growth and treatment

Tomato seeds were soaked in deionized water overnight in the dark. Four to five seeds have been transferred to polyethylene pots (tubes of diameter 43 mm; length 110 mm) containing 70 g air-dried soil (texture silt loam - USDA) sieved to 2 mm. The soil was brought to 80% water holding capacity (WHC) with distilled water and left for one week to equilibrate. Additionally, four wood sticks were used instead of plants to distinguish plant-mediated processes from abiotic ones. To avoid water stagnation at the bottom of the pots and ensure proper soil aeration, a layer of approx. 1 cm of perlite was placed under the soil. The underside of the pots was closed by a thin nylon membrane with 30 µm mesh. Aluminium foil was folded around the pots to prevent light contamination. The soil used for the experiment had the following properties: texture silt loam (USDA), WHC 37.2 mL 100 g^-1^; pH 7.8 (H_2_O), 7.4 (KCl); total carbonate 156 g CaCO_3_ kg^-1^; total organic carbon 13.52 g kg^-1^; total nitrogen 0.143 %; Olsen Phosphorus 57 mg kg^-1^, cation exchange capacity 16.9 cmol_(+)_ kg^-1^. Prior to the experiment, the soil was fertilized with the following salts (mg kg^-1^): NH_4_NO_3_ 500, KH_2_PO_4_ 250, KCl 70. The experiment was conducted in a climate chamber with the following environmental conditions: 14/10 h day/night, 24/19 °C, 70% relative humidity (RH) and 250 μmol m^-2^ s^-1^ light intensity. The pots were irrigated with MilliQ water every two days in order to maintain 80% WHC of the soil.

After 19 days of growing, once the plants fully developed the second leaf, uniform pots in terms of plant development (four plants per pot) have been selected and the plants foliarly sprayed with commercially available formulations of either the fungicide “Folicur” (1.125 kg ha^-1^, active ingredient: tebuconazole) or the insecticide “Confidor” (0.35 kg ha^-1^, active ingredient: *imidacloprid*). The fungicide tebuconazole and the insecticide imidacloprid are agrochemicals both registered for tomato defense/protection against powdery mildew and aphids-aleurodids, respectively. As control, some other plants were treated with either deionized water (C), the commercial herbicide “Round Up” (3.75 kg ha^-1^, active ingredient: glyphosate) or the N fertilizer urea (80 kg ha^-1^). As a further control, the aerial part of the woody sticks has been treated according to the experimental plan for the plants. The foliar applications foresee the spraying with the agrochemicals of the tomato shoots or the wood sticks until the whole aerial part was covered with a thin liquid film as described by labels’ recommendations. In all the treatments considered, the soil surface of plant and stick pots has been physically isolated from the above-ground applications by using plastic foils. Each agrochemical treatment included five replicates and the whole experiment was repeated independently three times.

### Sampling

Soil and plant samples were collected after two days from the agrochemicals’ application, which corresponds temporally to the onset of the *glyphosate*-induced toxicity symptoms in plants treated with this agrochemical. For the metabolomic analysis, the shoots, once separated from the roots at the collar and washed trice with deionised water, were then put immediately into liquid nitrogen to quench the metabolism and stored in the freezer (−80 °C) for later metabolite extraction. The rhizosphere soil, *i*.*e*. the soil adhering to the roots or the wood sticks, was carefully sampled by brushing and transferred into 2 mL Eppendorf tubes. Samples of bulk soil (unplanted pots) have been also considered. Sterile equipment was used for the whole process. The samples were stored in the freezer (−20 °C) for later DNA extraction.

### UHPLC-ESI/QTOF-MS untargeted metabolomics

The extraction of plant samples was carried out as previously set up (Lucini *et al*., 2018). Each replicate sample was comminuted with pestle and mortar in liquid nitrogen and an aliquot (1.0 g) extracted in 10 mL of 0.1% HCOOH in 80% methanol using an Ultra-turrax (Ika T25, Staufen, Germany), for 3 min in an ice bath. The extracts were then centrifuged at 10,000 x g for 10 min at 4 °C. Afterwards, the extracts were filtered using 0.22 µm cellulose syringe filters into amber glass vials for the further analysis.

Metabolomic profile, targeting non-volatile compounds, was determined through ultra-high-pressure liquid chromatography (UHPLC) coupled to quadrupole-time-of-flight hybrid mass spectrometry via an electrospray ionization source (UHPLC-ESI/QTOF-MS). A 1290 series liquid chromatograph, a JetStream dual electrospray and a G6550 mass spectrometer detector (all from Agilent Technologies, Santa Clara, CA, USA), were used. Chromatographic separation was achieved in reverse mode, on an Agilent Zorbax Extend-C18 column (75 × 2.1 mm i.d., 1.8 μm) and using a gradient elution (5% to 95% methanol, LCMS grade) over a 35 min run. The mass spectrometer was set to positive polarity SCAN mode (100–1000 m/z^+^ range), extended dynamic range mode, and 35.000 FWHM nominal resolution. Concerning electrospray conditions, nitrogen was used as drying gas (330 °C, 8 L min^-1^), nozzle voltage was 300 V and nebulizer pressure was 60 psi. Capillary voltage was 3.5 kV and lock masses (121.0509 and 922.0098 m/z^+^) were continuously infused during the whole chromatographic run.

Feature recursive extraction, together with mass and retention time alignment (0.1 min and 5 ppm respectively) were done in Profinder B.07 software (from Agilent Technologies). Annotation was done based on the ‘find-by-formula’ algorithm, using the combination of monoisotopic accurate mass, isotopes spacing and isotopes ratio. The database PlantCyc (www.pmn.plantcyc.org) was used for annotation purposes. Annotated compounds were filtered by frequency: only those compounds present in 75% of replications within at least one treatment were retained. Based on the strategy used, the identification process can be ascribed to Level 2 (putatively annotated compounds) as set out by the COSMOS Metabolomics Standards Initiative (http://cosmos-fp7.eu/msi.html).

The metabolomics dataset was interpreted in Mass Profiler Professional B.12.06 (Agilent technologies). Abundance was Log2 transformed and normalized at 75^th^ percentile and baselined against the median. Unsupervised hierarchical cluster analysis (Squared Euclidean distance, Ward’s linkage rule) was carried out using the fold-change based heat map to naively investigate relatedness among treatments. Thereafter, the raw metabolomic dataset was exported to software SIMCA 13 (Umetrics, Malmo, Sweden), pareto scaled and elaborated for orthogonal projection to latent structures discriminant analysis (OPLS-DA) as supervised interpretation. Once the model was available, outliers were investigated according to Hotelling’s T2 (95% and 99% confidence limits for suspect and strong outliers, respectively) and cross-validation carried out through CV-ANOVA (p < 0.001). Model parameters (goodness-of-fit R2Y and goodness-of-prediction Q2Y, were then recorded and permutation testing applied to exclude model overfitting (N = 300 – supplementary material). Finally, the variables importance in projection (VIP) approach was used for feature selection and Volcano Plot analysis was carried out by combining fold-change (FC – threshold: 2 folds) with ANOVA (P < 0.01, Bonferroni multiple testing correction) on discriminant compounds. The whole list of annotated compounds, as well as differential metabolites identified by Volcano Plot analysis, are provided as supplementary material.

### High-throughput sequencing of bacterial and fungal amplicons

The microbial ecology of bacteria and fungi was analyzed by Illumina high-throughput sequencing (HTS) of 16S and ITS amplicons respectively. The total microbial genomic DNA was purified from rhizosphere samples using the DNeasy PowerSoil Kit (QIAGEN GmbH, Hilden, Germany) following the manufacturer’s instructions. Bacterial and fungal diversity were analyzed targeting respectively the V3-V4 region of 16S ribosomal RNA (rRNA) and the Internal Transcribed Spacer 1 (ITS1) genomic region of ribosomal DNA (rDNA). Two-step PCR protocols were implemented in both cases to minimize amplification biases due to non-specific primer annealing, as previously described (Berry *et al*., 2011). The primers used to target the hypervariable V3 and V4 regions of bacterial 16S rRNA gene were 343F (5′-TACGGRAGGCAGCAG-3′) and 802R (5′-TACNVGGGTWTCTAATCC-3′). PCR amplification was carried out in a reaction mix containing 12.5 µL of Phusion Flash High-Fidelity Master Mix (Thermo Scientific Inc, Waltham, MA, U.S.A.), 1.25 µL of each primer (10µM concentration), 2 μL of DNA template (at 1ng/μL concentration), and 8 μL PCR ultrapure nuclease free water. Thermal cycling conditions were as follows: for the first step, an initial denaturation at 94 °C for 5 min, followed by 25 cycles at 94 °C for 30 s, 50 °C for 30 s, 72 °C for 30 s, followed by a final extension at 72 °C for 10 min. For the second step 2 µL of PCR products were amplified with an initial hold at 95 °C for 5 min, followed by 10 cycles of 95 °C for 30 s, 50 °C for 30 s, and 30 °C for 30 s; then, a final extension at 72 °C for 10 min. ITS region was amplified by using primers ITS-1 (5’-TCCGTAGGTGAACCTGCGG-3’) and ITS-2 (5’-GCTGCGTTCTTCATCGATGC-3’). PCR amplification was carried out with the reaction mix prepared using 12.5 μL of Phusion Flash High-Fidelity Master Mix, 1.25 μL of each primer (of 10 μM concentration), 2 μL of DNA template (of 1ng/μL concentration), and 8 μL of PCR ultrapure nuclease free water. Then, following thermal cycling conditions were implemented: for the first step, initial hold at 94 °C for 4 min, followed by 28 cycles of 94 °C for 30 s, 56 °C for 30 s, and 72 °C for 1 min; then, a final extension at 72 °C for 7 min. For the second step initial hold at 94 °C for 4 min, followed by 7 cycles of 94 °C for 30 s, 56 °C for 30 s, and 72 °C for 1 min; then, a final extension at 72 °C for 7 min. On the second steps for both fungal and bacterial amplicons, each sample was amplified using a dedicated forward primer with a 9-base extension at the 5’ end, which acts as a tag, in order to allow simultaneous analyses of all samples in a single sequencing run possible (Vasileiadis *et al*., 2012). PCR products generated from the second step were multiplexed as a single pool, for ITS and 16s amplicons separately, using equivalent molecular weights (20 ng). The pool was then purified using the solid phase reversible immobilization (SPRI) Agencourt AMPure XP kit (Beckman Coulter, Milan, Italy).

High-throughput sequencing data filtering, demultiplexing and preparation for concomitant statistical analyses were carried out as previously detailed (Spini *et al*., 2018). The first steps for sequences processing and filtering were the same for both 16S and ITS amplicons. Raw paired Illumina sequences were merged with the “pandaseq” script (Masella *et al*., 2012) with a minimum overlap of 30 bp between read pairs and 2 maximum allowed mismatches. Sequences were demultiplexed according to sample indexes and primers with the fastx-toolkit. Both bacterial and fungal amplicons were analyzed with taxonomy-based and OTU-based analyses: in the first case, all sequences were individually classified at taxonomical level against relevant database (GreenGenes for bacteria, UNITE for fungi), while in the OTU-based analyses, sequences were grouped at 97% similarities. For 16S amplicons, both operational taxonomic units (OTUs) and taxonomy-based matrixes were produced with a pipeline in Mothur (Schloss *et al*., 2009). For ITS amplicons, taxonomy-based analyses were also performed in Mothur, whereas OTUs were determined in UPARSE (Edgar, 2013).

For bacterial sequences, Mothur v.1.43.0 was applied in order to remove sequences with large homopolymers (≥10), sequences that did not align within the targeted V3–V4 region, chimeric sequences and sequences that were not classified as bacterial after alignment against the Mothur version of the RDP training data set. The resulting high-quality sequences were analyzed with Mothur and R following the OTU and the taxonomy-based approach. Sequences were first aligned against the SILVA reference aligned database for bacteria using the NAST algorithm and a kmer approach (Schloss, 2010) and then clustered at the 3% distance using the average linkage algorithm. OTUs were classified into taxa by alignment against the Greengenes database (McDonald *et al*., 2012).

ITS taxonomy-based analyses were conducted in Mothur: sequences shorter than 120 bp were discarded. We discarded homopolymers > 10 bp and chimeras, which were identified with the UCHIME algorithm implemented in Mothur, with the UNITE database version 6 as reference. The same database was used to classify the retained sequences and to eliminate non-fungal sequences. OTUs were produced in USEARCH with the –fastx_uniques and - cluster_otus commands. Sequences that did not belong to fungi were identified with the sintax command against the Utax reference database and discarded.

The OTU- and taxonomy-based matrixes obtained were analyzed in R to estimate the associated α and β diversity of the samples. The Good’s coverage estimate was calculated to assess the “percentage diversity” captured by sequencing. The most abundant OTUs identified were confirmed with BLAST (Basic Local Alignment Search) searches against the GenBank and the RDP database. Statistical analyses on OTU and taxonomy matrixes were performed in Mothur and R, to include hierarchical clustering with the average linkage algorithm at different taxonomic levels, and Principal component analysis (PCA) to assess the unconstrained samples grouping, Canonical correspondence analyses (CCA) to assess the significance of different treatments on the analyzed diversity. Metastats (Paulson *et al*., 2011) was applied to identify features that were significantly different between the treatments. Functional genes were inferred from 16S rRNA data using PICRUSt (Langille et al., 2013) and afterwards analyzed and visualized with STAMP (Parks et al., 2014).

The sequence data are publicly available in the National Center for Biotechnology Information (NCBI) database under BioProject accession number PRJNA642725.

## Results

### Effects on plant metabolism

Although no symptoms could be observed at phenotype level (except for *glyphosate*-treated plants), untargeted metabolomics highlighted that the treatments considered in this work imposed a profound biochemical reprogramming in tomato plants (Fig. 1A and 1B).

**Figure 1.**
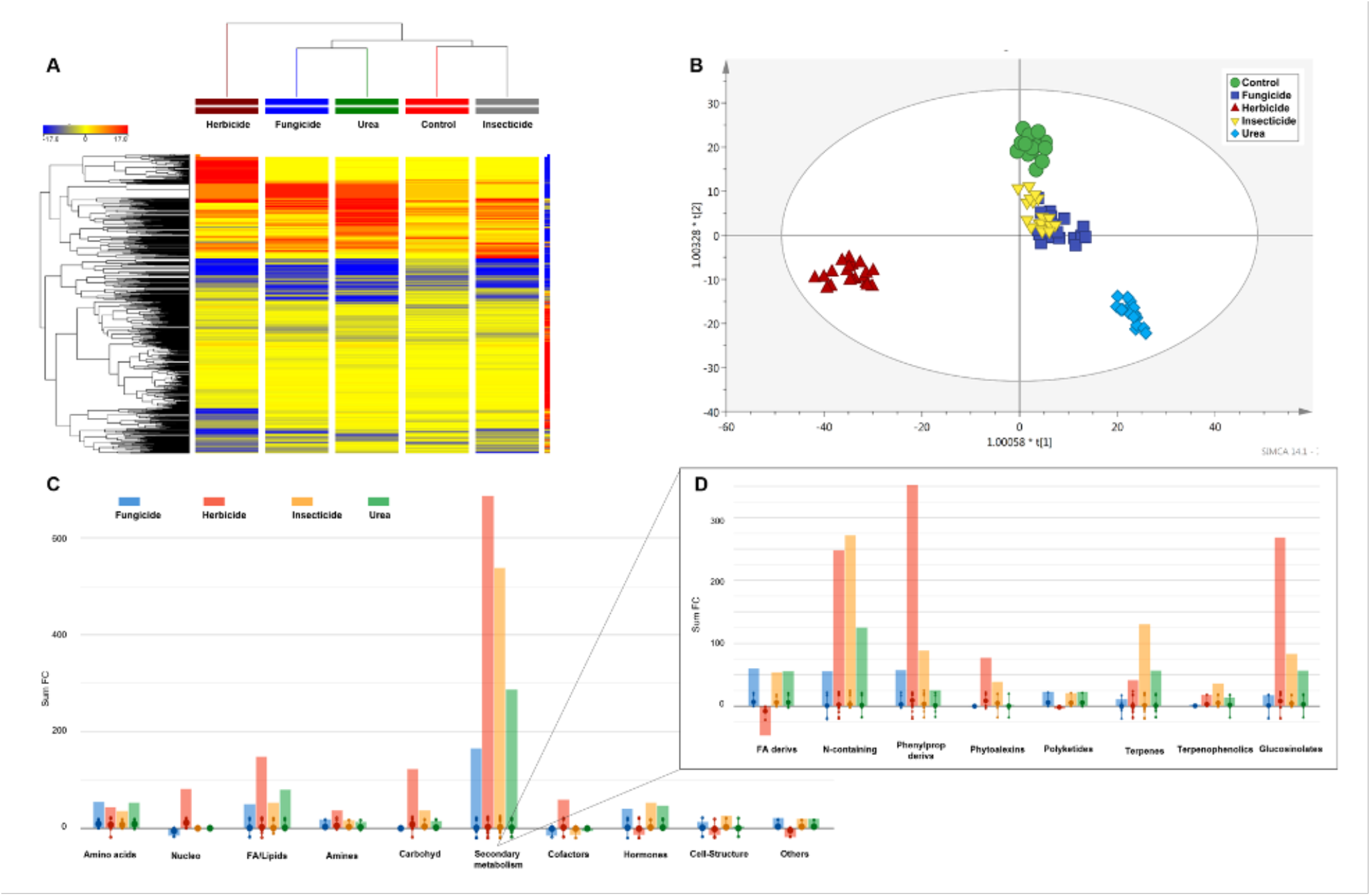
Multivariate statistics and interpretations from tomato metabolomic profile following treatment with the different agrochemicals: (A) unsupervised hierarchical clustering based on gold-change heatmaps (Euclidean distance, Ward’s linkage rule); (B) Orthogonal Projection to Latent Structures Discriminant Analysis (OPLS-DA) supervised modelling; (C) Biosynthetic processes being affected by the treatment (as provided from PlantCyc omic viewers output from differential metabolites) and (D) detail of secondary metabolites biosynthesis (expanded from 1C).

Unsupervised hierarchical clustering pointed out distinctive metabolomic responses following the application of the tested chemicals and suggested that such profound reprogramming might involve several other biochemical processes in addition to the detoxification pathways already known for xenobiotics. Supervised multivariate modeling confirmed the outcomes of hierarchical clustering, showing a marked alteration of metabolomic profiles and highlighting the treatment with *glyphosate* as the most distinctive profile (Fig. 1B). Notably, this supervised analysis suggested that the treatments with *imidacloprid* and *tebuconazole* resulted in rather comparable metabolomic fingerprints. Overall, more than 600 differential compounds were identified in tomato in response to the treatments and classified based on their functional class (Fig. 1C, 1D). As expected, secondary metabolism (biosynthesis of phenylpropanoids, glucosinolates, fatty acids derivatives, phytoalexins, and terpenes – Fig. 1D) underwent the most pronounced variations, together with lipids (mainly phospholipids and sterols), cofactors and phytohormones. Finally, the analysis of differential metabolites allowed identifying both the induction of reactive oxygen species and the consequent detoxification processes. Among the former, hydroxy-, oxo- and epoxy-derivatives of fatty acids, the peroxyl fatty acid 9-HPOTE, hexanoate, sphingosine, phosphocholine as well as mono- and diacylglycerides increased following pesticide treatments, whereas a decrease in the polyunsaturated fatty acid linolenate could be observed. Consistently, both quinols and quinones together with the previously mentioned glutathione exhibited evident down-accumulation trends. At the same time, 2-hexenal (involved in chloroplast detoxification of reactive carbonyls) and seleno-amino acids increased in treated plants.

### Effect on rhizosphere microbial community

The rhizosphere microbial community composition of tomato plants foliar sprayed with the different chemical products was assessed by means of Illumina high-Throughput Sequencing of phylogenetic markers for both bacteria (16S amplicons) and fungi (ITS). The rhizosphere samples of wood sticks treated similarly to tomato plants was used as control. The Operational Taxonomic Units from sequencing experiments were analyses by supervised Canonical Correspondence Analyses (CDA) and allowed identifying significant changes in the rhizosphere communities’ composition for both bacteria (Fig. 2A) and fungi (Fig. 2B). In more detail, the wood sticks grouped together (confirming a plant-mediated microbial shaping), whereas pesticide-treated plants differentiated from the control, confirming a distinctive and significant effect of each pesticide.

**Figure 2.**
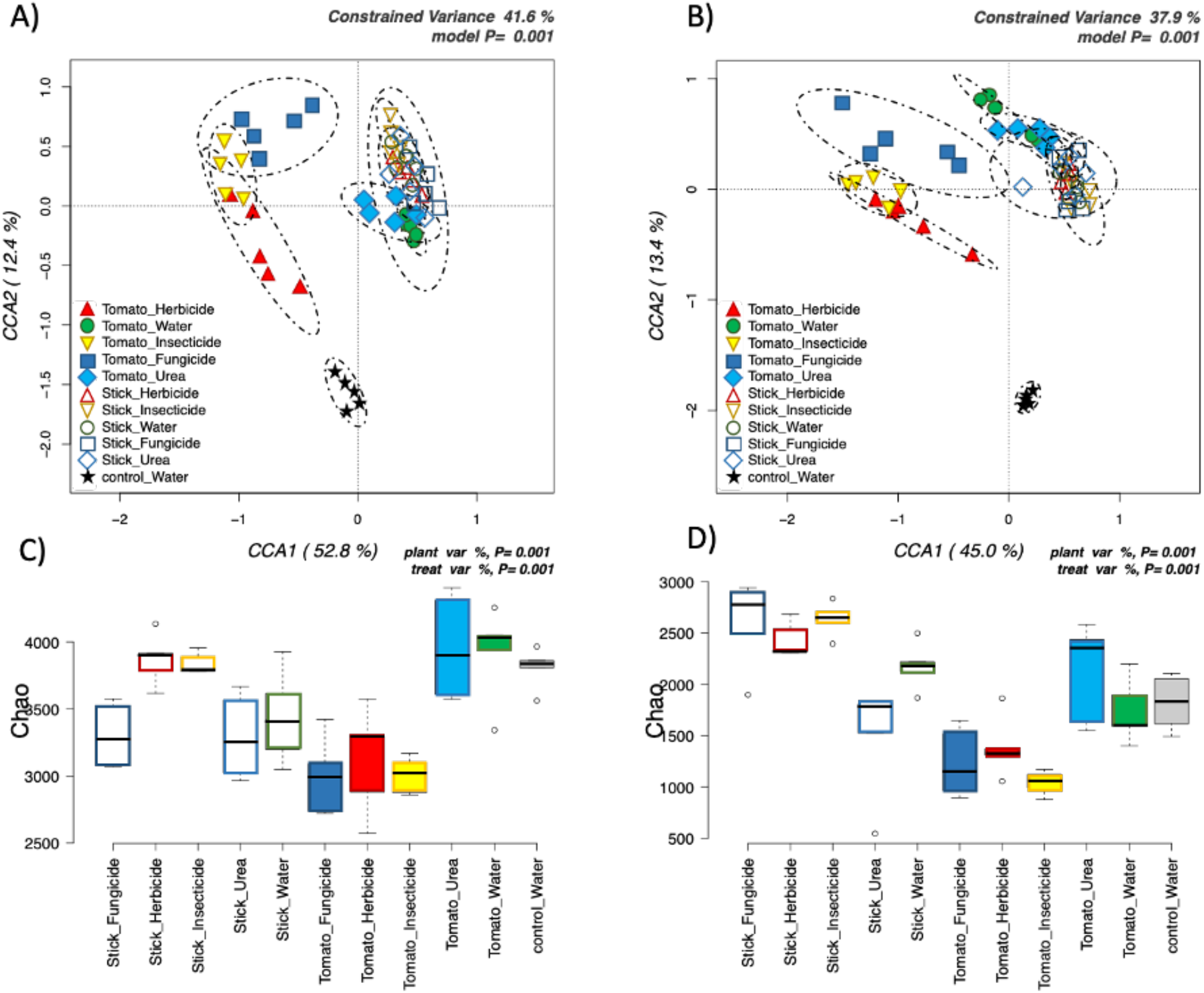
Canonical correspondence analyses (CCA) on the total OTU matrixes of bacteria (1a) and fungi (1b) testing the statistical significance of plant and treatment effects. Chao indexes of diversity were also calculated on the same matrixes for bacteria (1c) and fungi (d).

Looking at the biodiversity of these communities, both the total number of observed species (OTUs) per sample and the α-diversity Chao indexes indicate that the rhizosphere of pesticide-treated plants suffered from a significant decrease in the biodiversity of both bacteria (Fig. 2C) and fungi (Fig. 2D), compared to water- and urea-treated plants. These two latter treatments showed levels of diversity similar to those of the unplanted pots (bulk soil).

The OTUs covering all together 95% of total diversity were phylogenetically identified and compared. This further analysis showed significant differences among the treatments for bacteria and fungi (Fig. 3A and 3B). Indeed, some species significantly increased, and some others were significantly reduced by the pesticide treatments regardless of the agrochemical tested. The abundances of bacterial functional genes, inferred from 16S amplicons data indicate that the foliar application of the agrochemicals significantly increased several classes of functional genes such as cell motility, ions transports and metabolism, biosynthesis of secondary metabolites, degradation of different xenobiotics as well as cytochrome P450 mediated detoxification mechanisms (supplementary material).

**Figure 3.**
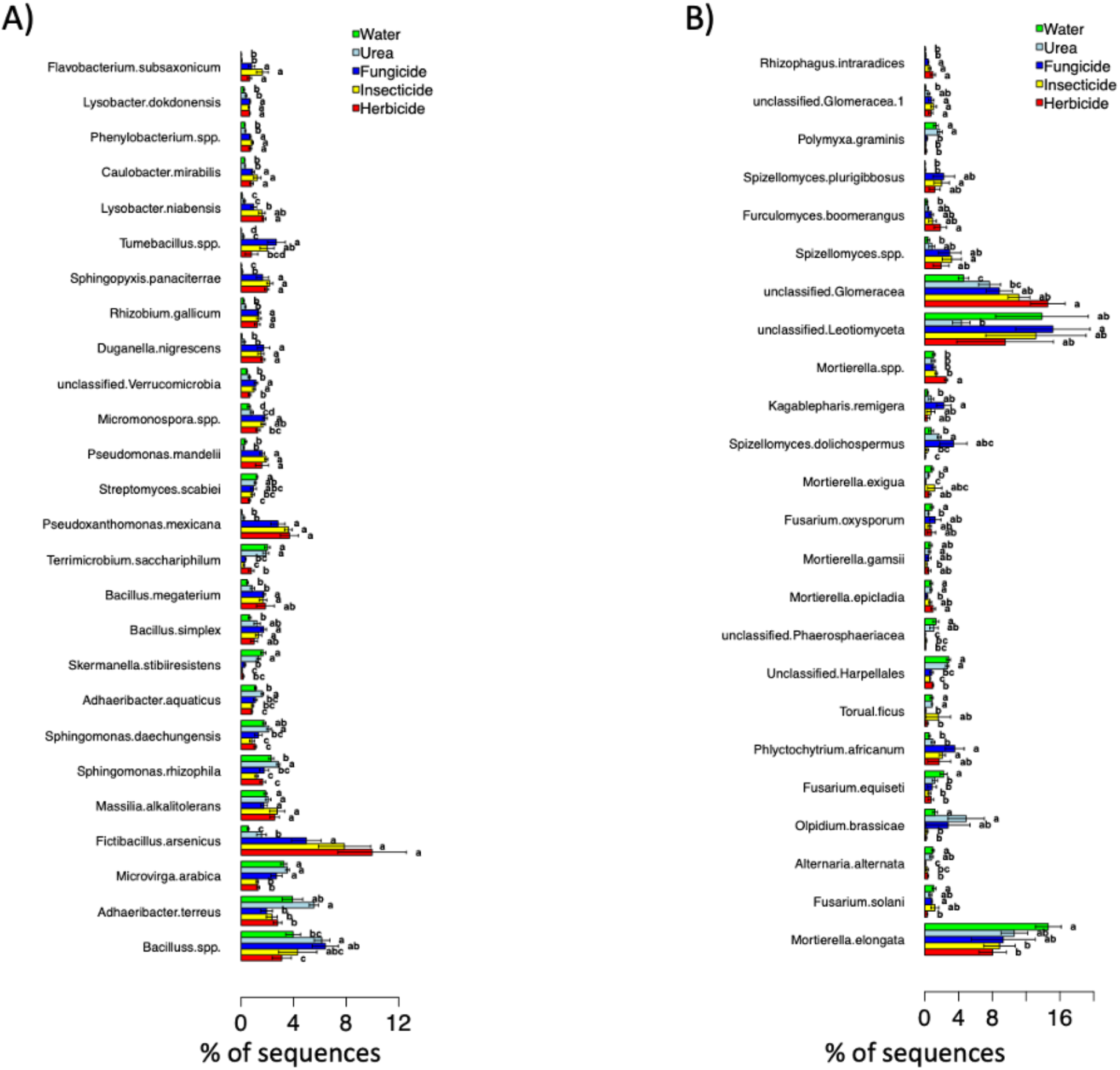
Metastats analysis testing significant differences in bacteria (a) and fungi (b) between treatments for tomato samples. OTUs representing the 95% of total diversity are reported, and classified at the lowest taxonomical level, in most cases the species level.

## Discussion

### The concealed metabolic reprogramming

The occurrence of some changes at metabolite level is in agreement with previous findings (Parween *et al*., 2016; Sun *et al*., 2018). It is well known the pivotal role played by glutathione in plant response to xenobiotics, likely acting on both redox imbalance and detoxification *via* conjugation. The concurrent accumulation of phenylpropanoids is considered to be a component of the radical scavenging strategy carried out by plants.

Notably, the agrochemicals’ treatments considered in the present work also altered the phytohormone profile of the tomato plants. This aspect is particularly crucial because it opens the possibility of a wide range of morphological, growth, and productive differences in the plants. In particular, the stimulation of jasmonates biosynthesis is coherent with the stress condition postulated above. In fact, the signaling cascade of this hormone affects several processes such as interaction with pathogens and beneficial organisms, response to abiotic stresses, root growth, flowering, and light perception (Wasternack and Hause, 2013). However, gibberellins and cytokinins were also affected by the treatments. These two hormones are a part of the whole multilevel hormone crosstalk that finally affects plant growth and response to stimuli. Moreover, they act antagonistically in leaf formation, epidermal differentiation, and meristem maintenance (Gan *et al*., 2007). Finally, distinctive signatures could also be gained for the brassinosteroids, with very distinctive responses according to the treatment considered. These polyhydroxylated steroidal plant hormones are relatively less studied. They exhibit growth-promoting activity and are involved in several processes, including cell elongation, seed germination (Haubrick and Assmann, 2006), stomata control and response to drought (Daszkowska-Golec and Szarejko, 2013), as well as in response to abiotic stresses such as salinity or heavy metals (Javid *et al*., 2011; Bartwal *et al*., 2013). Interestingly, brassinosteroids are strictly connected to the production of reactive oxygen species (Xia *et al*., 2010) and their role in alleviating pesticide-related oxidative stress has been postulated (Shahzad *et al*., 2018).

Plants can adopt a metabolic cascade of enzymatic reactions for detoxifying and storing xenobiotics that generally involves three distinct phases, *i*.*e*., i) activation, ii) true detoxification and iii) excretion (Coleman *et al*., 1997). While activation mainly involves P-450 monooxygenases and peroxidases, glutathione- and glucosyl-transferases conjugation is the typical enzymatic reaction of the true detoxification phase, commonly resulting in a reduction of toxicity to plant (Schröder *et al*., 2001). Furthermore, trafficking mechanisms like transport and storage into the vacuole rather than cell wall binding and excretion are also essential steps in the overall detoxification process (Theodoulou, 2000).

Altered metabolic functions of plants following pesticide treatments have already been reported in several works (Verkleij *et al*., 2009; Parween *et al*., 2016; Sun *et al*., 2018). Compared to a mere detoxification process, these metabolic alterations seemed to be much more pronounced and severe. In fact, they often resulted in changes at the morphological level of the whole plant (*e*.*g*., height, number of branches, etc.), production of reactive oxygen species, accumulation of osmolytes, membrane integrity alteration and/or malfunction of the carbon and nitrogen metabolism (Parween *et al*., 2011, 2012, 2016; Sun *et al*., 2018). It is interesting to note that these previous clues are perfectly in agreement with the distinct and profound changes at the metabolome level detected in our work. In this regard, it is worth noting that the three pesticides here used (either *tebuconazole, imidacloprid* or the positive control *glyphosate*, sprayed onto leaves) are very different from a chemical point of view and are known to target somewhat distant metabolic processes in the pests or weeds. Nonetheless, the modulation of plant metabolism in tomato plants was extensive and accounted for both common and specific responses.

### Shift in microbial community at rhizosphere level

The large majority of the cases, the effects on the microbial communities caused by chemical inputs include a change in biodiversity and community structure, together with biomass abundance (Puglisi, 2017). Considering the agrochemicals analyzed in our work, *tebuconazole* and *imidacloprid* are reported to cause a decrease of the microbial diversity and activity (Muñoz-Leoz *et al*., 2011; Cycoń and Piotrowska-Seget, 2015; Mahapatra *et al*., 2017; Storck *et al*., 2018), while no changes or only transient modifications in the soil microbial communities have been reported for *glyphosate* (Weaver *et al*., 2007; Mijangos *et al*., 2009; Dennis *et al*., 2018). It is interesting to note that, so far, the scientific interest has been focused on the direct soil contamination by agrochemicals and their consequences from an agronomical and/or environmental point of view. For this reason, the question of a possible plant-mediated effect of agrochemicals on the rhizosphere microbial component has remained open until now. In this respect, our results show for the first time that chemical compounds like agrochemicals can actually shape the rhizosphere microbiome *via* the physiological responses to these products foliar applied.

With respect to bacteria, an increase of *Pseudoxanthomonas Mexicana* (Nayak *et al*., 2009), usually described as a xenobiotic degrader, has been observed as a consequence of the treatments. Interestingly, the plant-mediated responses to agrochemicals also caused the increase of species featuring plant-growth promoting traits such as *Pseduomonas mandelii* and *Bacillus simplex* (both species with fungal-disease suppression abilities (Mavrodi *et al*., 2012; Schwartz *et al*., 2013), *Rhizobium gallicum* (a diazotrophic nodulating symbiont (Amarger *et al*., 1997) and *Bacillus megaterium* (related to biostimulant properties (Zou *et al*., 2010), resistance inducing (Chakraborty *et al*., 2006) and also biodegradation activities towards several agrochemicals (Bhadbhade *et al*., 2002; Yao *et al*., 2011; Liu *et al*., 2014). Consistently, the inference of the relative abundance of functional genes indicates a significant increase of species with xenobiotic-degrading abilities and enrichments in species with plant growth-promoting activities. It is interesting to highlight the increase in cytochrome P450 systems, which indicates the activation at the bacterial level of the detoxification processes of xenobiotics *via* oxidative pathways (Hannemann *et al*., 2007).

With respect to the fungal community, the foliar application of two of the three agrochemicals (*tebuconazole* and *glyphosate*) caused a significant restraint of the *Mortierella elongata* abundance, a fungus with plant-growth promoting and xenobiotic degrading activities (Li *et al*., 2018). Besides, two other species of the same genus, *i*.*e*., *Mortierella exigua* and *Mortierella gamsii*, were also restrained in their presence by *tebuconazole* or *imidacloprid*, respectively. Nevertheless, no precise ecological role can be defined for these two species. Another relevant effect connected to the treatments is the significant increase of *Rhizophagus intraradices*, an arbuscular mycorrhizal fungus with the ability to reduce the drought stress in plants (Li *et al*., 2014) and of unclassified Glomeracea, a family comprising several arbuscular mycorrhizal species.

Overall, it is possible to argue that our results on microbial communities of the rhizosphere reveal two complementary effects as a consequence of the plant treatments considered. In fact, while a reduction in biodiversity has been recorded, on the other hand there is a clear modulation of the bacterial and fungal species featuring important functional roles. Indeed, these results highlight a remodulation of the rhizosphere microbial diversity where microbial groups that can help the plant under chemical-stress (PGPR, xenobiotic degraders), are enriched through exudates-mediated communication systems.

### Overall considerations on the hidden effects caused by agrochemicals

To date, the pesticide-related indirect effects mediated by the crop have not yet been taken into consideration and therefore are still hidden, despite their possible environmental and production consequences. Results here reported clearly show that systemic pesticides foliar sprayed to tomato plants to protect them against pests, are recognized by the plant as xenobiotics and then subjected to a metabolic process of deactivation. The induction of this type of response has unavoidably energetic and metabolic costs for the plant, which causes thus a profound alteration of the metabolites profile in the plant itself. Regardless of the specific processes affected by the pesticides used in the present study, the hidden effects induced by the treatments were profound and involved critical aspects of the plant metabolism, potentially impacting growth and yields. Interestingly, the changes occurring at plant metabolome open the possibility that the nutritional, functional, and commercial quality traits of the products might be affected. From a food production perspective, the reorientation of the crop metabolism can induce modulation a whole series of metabolites, including those having functional significance. Therefore, the knowledge of these hidden aspects of a pesticide could be strategic in the framework of modern and sustainable agriculture but also has obvious implications in terms of food safety and food quality.

Furthermore, considering that roots release in the soil several exudates in a controlled manner aimed at interacting with neighboring organisms, a link between ongoing metabolic processes and exudation patterns has been postulated (Chaparro *et al*., 2013; Vranova *et al*., 2013; Canarini *et al*., 2016). In the present work, we postulated that the metabolic reprogramming of plants induced by the pesticide applications, affecting the metabolites’ pool available for the root exudation process, is very likely to reshape the microbial community in the rhizosphere.

Concerning the reciprocal plant-microorganism-soil interactions in the growth system, there is no doubt that the process of root exudation in the rhizosphere, fundamental for specific recruitment and consequent colonization of the root tissues by microorganisms, is strongly quanti-qualitatively influenced by the variability of the available metabolites. In our study, both a considerable reorganization of the plant’s metabolism and a different microbial ecology of the rhizosphere has been recorded after foliar application of the pesticides, indicating a close relationship between the two phenomena. This hypothesis is further corroborated by the pesticide-specific pattern. In this respect, it is interesting to note that the nutrient cycles and nutrients availability in the rhizosphere are the results of a series of specific interactions between these three components (root, soil, microorganisms) whose extent is also determined by the nature and the quantity of the released exudates. However, in shaping the rhizosphere, the advantage arising from the stimulation of plant-growth promoting bacteria should not be overlooked.

In conclusion, the setup of the most appropriate practice of pesticide-based defense of crops at the field scale cannot ignore these still unknown phenomena. Nowadays, it is impossible to imagine the cultivation of crops without the application of agrochemicals, and the knowledge of these hidden aspects related to pesticide use becomes fundamental. This information is essential to identify any contraindications/restrictions in some of their uses/applications, as occurs for medicines in the human field. However, in the framework of sustainability in agriculture, both possible limitations and positive aspects (directly in terms of crop defense and indirectly in terms of a shift in microbial communities) worth to be taken into account.

## Supplementary data

**Supplementary Table 1:** Dataset of metabolites annotated by untargeted metabolomics in plant samples following foliar application of different agrochemicals.

**Supplementary Figure 1:** Box-plots showing the relative abundance of the predictive functionality at rhizosphere level, across the different treatments, as investigated via PICRUSt.

## Acknowledgements

The authors are grateful to the “Romeo ed Enrica Invernizzi” foundation (Milan, Italy), for the kind support to the metabolomics facility at Università Cattolica del Sacro Cuore. This work is the result of a postdoctoral contract for the training and improvement abroad of research staff (Begoña Miras-Moreno; 21252/PD/19) financed by the Consejería de Empleo, Universidades, Empresa y Medio Ambiente of the CARM, through the Fundación Séneca-Agencia de Ciencia y Tecnología de la Región de Murcia (Spain).

## Author contributions

BMM, GS, ET, RT and MSZ carried out experiments and analyses. LL, BMM, LB, TM, YP and EP analysed and interpreted the data. LL, TM, YP and TM drafted the manuscript. SC and MT supervised the work and critically revised the draft manuscript. All authors approved the final version.

## Data availability statement

Metabolomics data are provided as supplementary material. The sequence data are publicly available in the National Center for Biotechnology Information (NCBI) database under BioProject accession number PRJNA642725.

## Abbreviations

BLAST: Basic Local Alignment Search)
CDA: Canonical Correspondence Analyses
ITS1: Internal Transcribed Spacer 1
NCBI: National Center For Biotechnology Information
OPLS-DA: Orthogonal Projection to Latent Structures Discriminant Analysis
OTUs: Operational Taxonomic Units
PCA: Principal Component Analysis
rDNA: ribosomal DNA
rRNA: ribosomal RNA
SPRI: Solid Phase Reversible Immobilization
UHPLC: Ultra-High-Pressure Liquid Chromatography
UHPLC-ESI/QTOF-MS: Quadrupole-Time-of-Flight Hybrid Mass Spectrometry
VIP: Variables Importance in Projection
WHC: Water Holding Capacity

## Notes

### Competing Interest Statement

The authors have declared no competing interest.

